# Aged Breast Extracellular Matrix Drives Mammary Epithelial Cells to an Invasive and Cancer-Like Phenotype

**DOI:** 10.1101/2020.09.30.320960

**Authors:** Gokhan Bahcecioglu, Xiaoshan Yue, Erin Howe, Ian Guldner, M. Sharon Stack, Harikrishna Nakshatri, Siyuan Zhang, Pinar Zorlutuna

## Abstract

Age is a major risk factor for cancer. While the importance of age related genetic alterations in cells on cancer progression is well documented, the effect of aging extracellular matrix (ECM) has been overlooked. Here, we show for the first time that the aging breast ECM is sufficient to drive normal mammary epithelial cells (KTB21) to a more invasive and cancer-like phenotype, while promoting motility and invasiveness in MDA-MB-231 cells. E-cadherin membrane localization was lost in KTB21 cells cultured on the decellularized breast matrix from aged mice. Cell motility, cell invasion, and inflammatory cytokine and cancer-related protein production were increased significantly on the aged matrix, and many genes related to invasion were upregulated. Strikingly, we showed using single cell RNA sequencing that the aged matrix led to enrichment of a subpopulation of KTB21 cells that highly expressed epithelial-mesenchymal transition (EMT) and invasion-related genes. Lysyl oxidase (*LOX*) knockdown reverted the aged matrix-induced changes to the young levels; *LOX* siRNA treatment prevented the loss of E-cadherin membrane localization, and reduced cell motility, cell invasion, and cytokine and cancer-related protein production. Finally, we showed that the biophysical, mechanical and biochemical properties of the breast ECM were altered dramatically upon aging. Analyzing these factors and studying the differential response of the epithelial cells to young and aged ECMs could lead to identification of new targets for cancer treatment and could pave the way for the discovery of new therapeutic options.

## 1. Introduction

Cancer incidence increases dramatically with age^1^, suggesting that aging may promote tumorigenesis. This clinical observation has so far been attributed to effects of aging on the genetic makeup of cells^2–4^, and therefore aging-associated changes in tissues have conventionally been investigated at the cell level^5–8^. While the effect of aging-associated dysregulation of the cellular machinery on cancer initiation is well documented, the effect of aging-associated changes in the microenvironment, specifically the extracellular matrix (ECM), is overlooked. Neoplastic transformation of rat liver epithelial cells leads to higher rates of tumor formation when cells are transplanted into aged rat livers compared to young^9^, suggesting that the aged microenvironment plays important roles in tumor initiation and progression. However, whether and how the aged ECM contributes to cancer initiation and progression is not known.

Even slight differences in the biochemical composition, stiffness, and structure of the ECM may lead to a significant difference in cellular response^10–13^. For instance, while collagen I promotes epithelial-mesenchymal transition (EMT)^14,15^, collagen XV prevents it^15^. In the aged microenvironment, collagen density decreases due to loss of ECM integrity, leading to a greater invasive ability of tumor cells^1,16^. Decrease in fiber thickness is another age-related alteration in the ECM that could be contributing to metastasis^17^. On the other hand, substrate stiffness leads to reprogramming of normal mammary epithelial cells into tumor cells^18^ and cancer cell-mediated blockade of adipocyte differentiation and maturation^19^. Culture of malignant progenitors on soft substrates reverts them to normal epithelial cells^20^. However, how the ECM changes with aging, and these changes influence the normal epithelial cell behavior have not been investigated.

Despite the fact that breast cancer is one of the most common and widely studied cancer types, there is a dearth of research on age-related alterations in the ECM of the mammary gland. Here, we provide a full evaluation of the structural, mechanical, and biochemical changes that occur in the mouse breast ECM upon aging, and investigate the effect of aged decellularized breast matrices on the normal and cancerous human mammary epithelial cells. This is the first study to report the aging-associated changes in the breast ECM and investigate the exclusive effect of aged ECM on EMT and invasiveness of both normal and cancer cells.

## 2. Results

### 2.1. Aging leads to thicker collagen fibers, greater modulus, and altered biochemical composition in the breast

To understand the age-related changes in ECM properties of the breast, we characterized tissues from young (3-6 months old) and aged (22-25 months old) mice before (native tissue) and after decellularization (decellularized matrix, decell) and delipidization (delipidized matrix, delip). It is known that breast becomes less dense with aging due to increased fat content^21,22^. Here, we observed lumpy, patchy, and dense collagen network, and wavier, more twisted, and thinner (~30% decrease) collagen fibers in the aged tissue, regardless of the decellularization/delipidation status (Figs. 1a, 1b, and S1a).

**Fig. 1.**
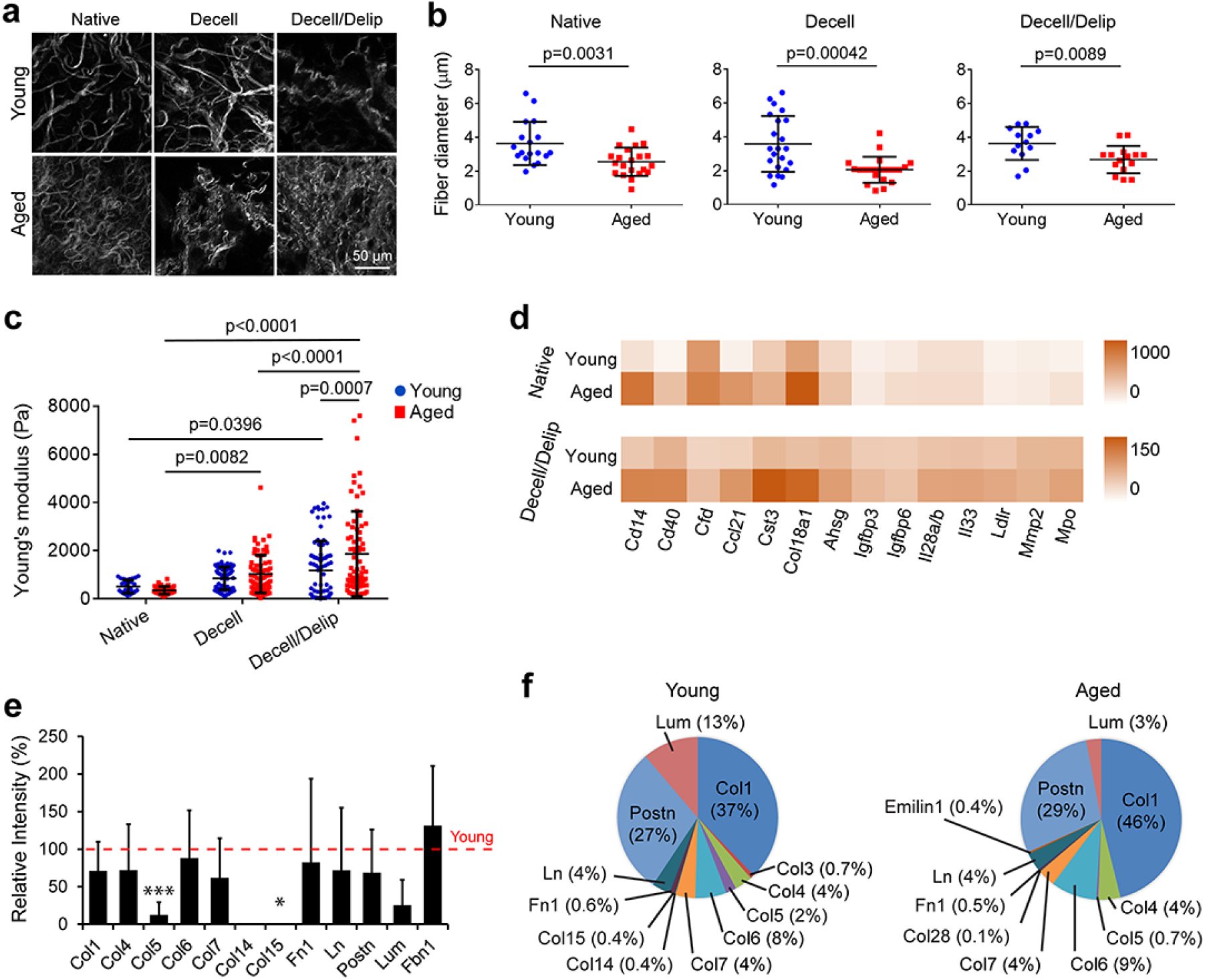
Aging leads to thinner collagen fibers, denser collagen mesh, increased stiffness and cytokine levels, and decreased level of structural proteins in the breast tissue. (a) Second harmonic generation (SHG) imaging in multiphoton microscopy showing the collagen fiber structure in the matrices. (b) Fiber diameter as quantified from the SHG images in (a). n= 18 for the young and 21 for the aged native tissues. n=22 for the young and 21 for the aged decellularized matrices. n=13 for the young and 15 for the aged delipidized matrices. (c) Elastic modulus of the breast tissues before (native) and after removal of cells and lipids as determined with nanoindentation testing. n=5 tissues/group. (d) Cytokine profiling of the native breasts and decellularized matrices as quantified by chemiluminescence-based immunoassay. n=3 pooled samples. Results representative of two independent experiments. Also see Table S1. (e, f) Mass spectrometric analysis showing the most abundant proteins in the matrices. (e) Signal intensity of the proteins in the aged matrix relative to that in the young (red line). *p<0.05, and ***p<0.001. (f) Percentages of proteins present in the matrices. n=3 for young and 5 for aged matrices. Data are presented as the mean ± SD. Student’s t-test was applied for (b) and (e), and one-way ANOVA followed by Tukey’s post hoc for (c).

Stiffness changes with age in various tissues^23^; however, the effect of aging on mechanical properties of the breast has not been studied, although ultrasound elastography based measurements on patients have indicated a slight increase in stiffness of the echogenic homogeneous fibroglandular tissues in breast upon aging^22^. Here, using nanoindentation testing we found that the elastic modulus of the native breast tissue decreased with age (young: 509 ± 275 Pa, and aged: 356 ± 162 Pa) (Fig. 1c). However, the modulus of the aged matrix (1867 ± 1765 Pa) was greater than the young (1180 ± 1226 Pa) after decellularization/delipidation (p<0.0089). Decellularization/delipidation led to higher modulus, and this increase was more pronounced on the aged matrices (p<0.0001, decell/delip matrices compared to native) compared to the young (p=0.04, decell/delip matrices compared to native), suggesting that aged tissues contained more fatty components, yet the fibrous component of their ECM is stiffer.

The effect of aging on the matrisome of the mammary gland has not been well-studied, but cytokine levels increase with age in other tissues^1,24^, while collagen I and IV, laminin 1, and periostin decrease^25–27^. Our data show that the levels of most proteins, especially Cd14, Cd40, Ccl21, cystatin C (Cst3), endostatin (Col18a1), fetuin A (Ahsg), and myeloperoxidase (Mpo), were elevated with age, and this trend was preserved after decellularization/delipidation, although total protein levels were lower (Figs. 1d and S1b, and Table S1).

Conversely, the amount of structural ECM proteins was reduced with age (except for fibrillin 1 (Fbn1), which increased slightly), and particularly remarkable were the decreases in collagens V (p=0.001) and XV (p=0.044) (Fig. 1e). On the other hand, collagen I (Col1) constituted the majority (37-46%) of the detectable proteins, and its proportion in the proteome increased with age, although its amount was lowered (Figs. 1f and S1c). The proportion of Lumican (Lum) decreased with age, while those of other proteins stayed relatively stable. Hematoxylin and eosin staining of the native tissues showed a reduction in the epithelium portion in the aged tissue as expected (Fig. S1d). Collectively, these data show that structural proteins decrease with age, while the levels of most cytokines and glycoproteins such as Fbn1 were elevated.

### 2.2. Aged microenvironment leads to EMT-like and invasive cell behavior

Next, we examined the effect of aging microenvironment on the behavior of KTB21 human mammary basal epithelial cells, which we established previously^28^. For testing the spheroid forming and retaining ability as well as viability of KTB21 cells we used Matrigel-coated decell/delip matrices to induce spheroid formation, whereas for the migration and invasion studies we used Matrigel-free matrices. KTB21 cells formed spheroids on the Matrigel-coated young and aged matrices (Fig. 2a). Interestingly, while spheroids remained intact on the young matrices, they started to deform on the aged matrices at day 10 of incubation and disappeared by day 15, evident from bright field and second harmonic generation (SHG) micrographs (Fig. 2b). Furthermore, the spheroids on the aged matrices had lower circularity (p<0.0001) and greater aspect ratio than those on the young (p<0.005). Cell viability was high (~85%) on both matrices (Fig. 2a), showing that deformation of the spheroids was not due to cell death.

**Fig 2.**
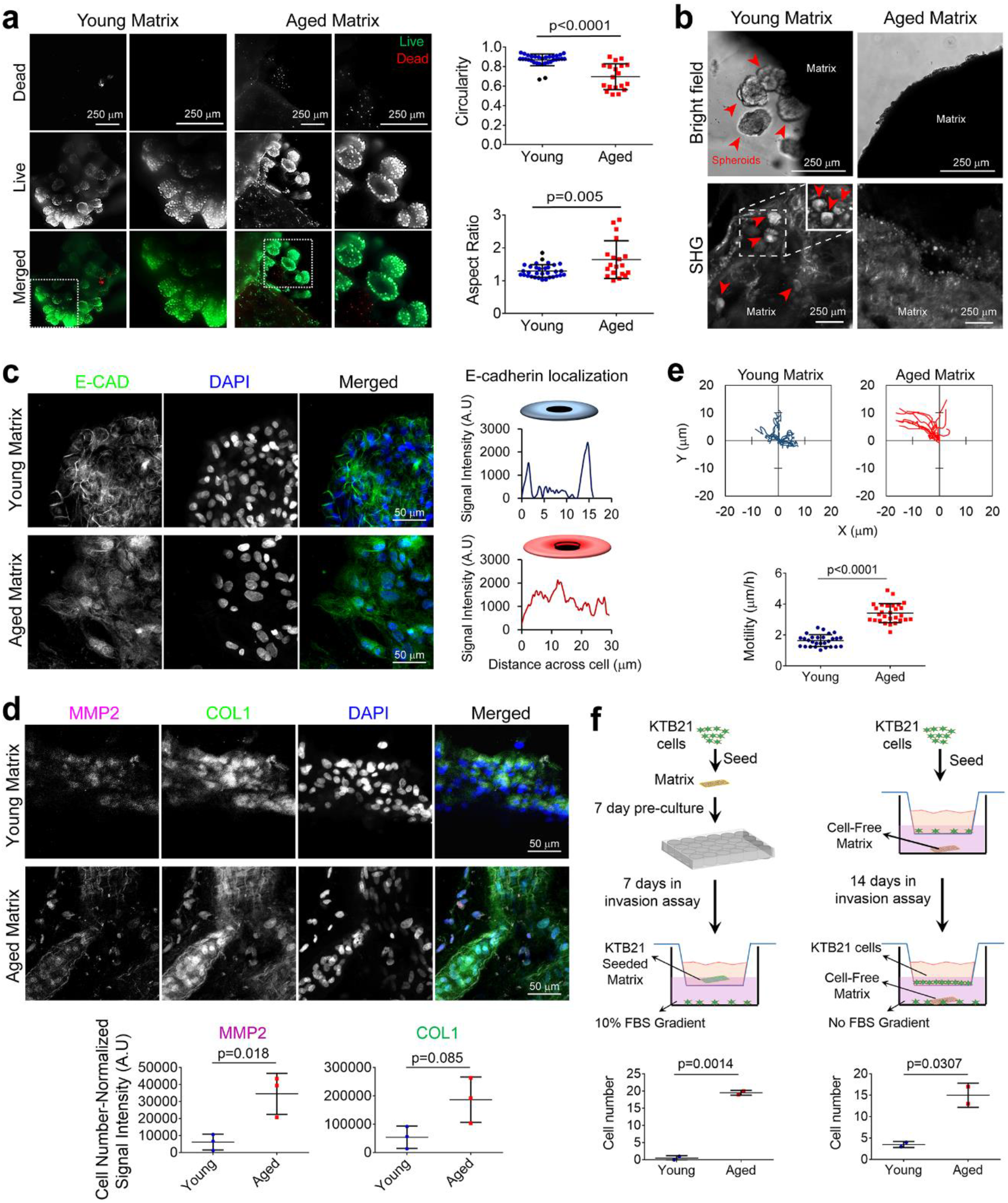
Aged breast ECM leads to deformation of KTB21 cell spheroids, delocalization of E-cadherin from membrane, and increase in motility and invasion. (a) Live-dead stained KTB21 spheroids on mouse breast ECMs (day 14). Left: fluorescence microscopy images showing viability of cells in spheroids. Dashed squares: magnified regions. Right: quantification of the circularity and aspect ratio of the spheroids. Green: live (calcein-AM), red: dead cells (ethidium homodimer-1). (b) Cell spheroids on the matrices (day 15). Top: bright field microscopy images. Bottom: second harmonic generation (SHG) microscopy images. Inset: image at a deeper focal plane of the dashed square.

Arrowheads show spheroids. (c, d) E-CAD, MMP2, and COL1 staining (day 15). (c) E-CAD localization in cells. Left: representative fluorescence microscopy images of matrices. n=3. Right: plot profile of E-CAD signal intensity across a representative cell. n=6 cells/matrix for young and 9 for aged. (d) MMP2 and COL1 staining. Top: fluorescence microscopy images of matrices. Bottom: quantification of the signal intensities. n=3. (e) Cell migration on matrices. Time-lapse images taken at 15 min intervals for 3 h. Top: cell trajectories. Bottom: motility. (f) Cell invasion through transwell inserts. Left: invaded cell number against FBS gradient. n=2 matrices. Right: invaded cell number towards decell/delip matrices used as chemoattractants in the bottom chamber. n=2 matrices. E-CAD and COL1: Alexa fluor 488 (green), MMP2: Alexa fluor 647 (magenta). Nucleic acid: DAPI (blue). Quantifications are performed using the Fiji software. Data are presented as the mean ± SD. Statistical tests: two-tailed student’s t-test.

The disappearance of spheroids on the aged matrices could be due to reduced cell-cell interactions or increased degradation of the matrix. Thus, at day 15 of culture we stained the KTB21 cells for E-cadherin (E-CAD), matrix metalloproteinase 2 (MMP2), and COL1 expression. Strikingly, while E-CAD was localized to cell membrane on the young matrix, no or little membrane localization was observed on the aged matrix (Figs. 2c and S2a). Additionally, higher amounts of MMP2 (p=0.018) and COL1 (p=0.085) were deposited on the aged matrices (Fig. 2d and S2b).

We then investigated the migration and invasion behavior of KTB21 cells on the matrices. Remarkably, cell motility was higher on the aged matrix than the young (p<0.0001) (Fig. 2e). For invasion, we used two approaches. In the first, we pre-incubated the cells on the matrices for 7 days, followed by a 7-day incubation in transwell inserts (upper chamber) against a 10% FBS gradient (bottom well) (Fig. 2f, left). The number of cells that invaded through the inserts was higher in the presence of aged matrix than young (p=0.0014). In the second approach, we seeded KTB21 cells in transwell inserts and placed the decell/delip matrices in the bottom wells (assuming they would release cytokines as chemoattractants) (Fig. 2f, right). After 14-day incubation, the number of cells that invaded towards the aged matrix was significantly higher than that of cells invaded towards the young (p=0.0307), indicating that cytokines released from the aged matrices promote the invasion of cells.

Similar migratory and invasive behavior was observed with the MDA-MB-231 breast cancer cells. Cell motility on the aged matrix was greater than that on the young (p<0.0001) (Figs. S3a and S3b). Interestingly, cells migrating on the aged matrix were mainly spindle-like (smaller circularity (p<0.0001) and roundness (p=0.003)), whereas those on the young matrix were round (Fig. S3c). Invading cell count was also higher in the presence of aged matrix (Fig. S3d).

### 2.3. Aging microenvironment leads to upregulation of cancer-associated genes and enrichment of a cell cluster defined by EMT transcriptome

To dissect the phenotypic changes we observed in normal epithelial cells, we performed single-cell RNA-sequencing to investigate how 20-day culture on aged matrix impacts the transcriptome. Dimension reduction using t-distributed stochastic neighbor embedding (t-SNE) showed that KTB21 cells maintained their basal phenotype, with a large proportion (70%) of the cells expressing the basal epithelial cell markers, *KRT5* and *KRT14*, and a smaller subset expressing the luminal markers, *KRT8* and *KRT18* (Fig. S4a), in line with previous characterization^28^. Examination of bulk cells showed similar transcriptomes on the young and aged matrices (Fig. S4b). However, 29 genes were significantly upregulated in cells on the aged matrix (adjusted p<0.05), including *LOX, LOXL2, MME, GJA1, MALAT1, NEAT1, TIMP3, IGFBP3, and SERPINE1* (Fig. 3a). We then performed unbiased clustering to examine the transcriptomic heterogeneity induced by culture on the matrices. We identified 9 transcriptional clusters (Fig. 3b). Interestingly, although cells cultured on young and aged matrices were represented in all 9 clusters, cells from aged matrices were enriched in clusters 0 (young – 164 cells, aged – 275 cells) and 7 (young – 44 cells, aged – 123 cells), which were transcriptionally close to each other, but divergent from the majority of the cells. Most of the genes that were upregulated in the aged microenvironment were also highly expressed in cluster 7 cells compared to cells in other clusters (Fig. 3c).

**Fig. 3.**
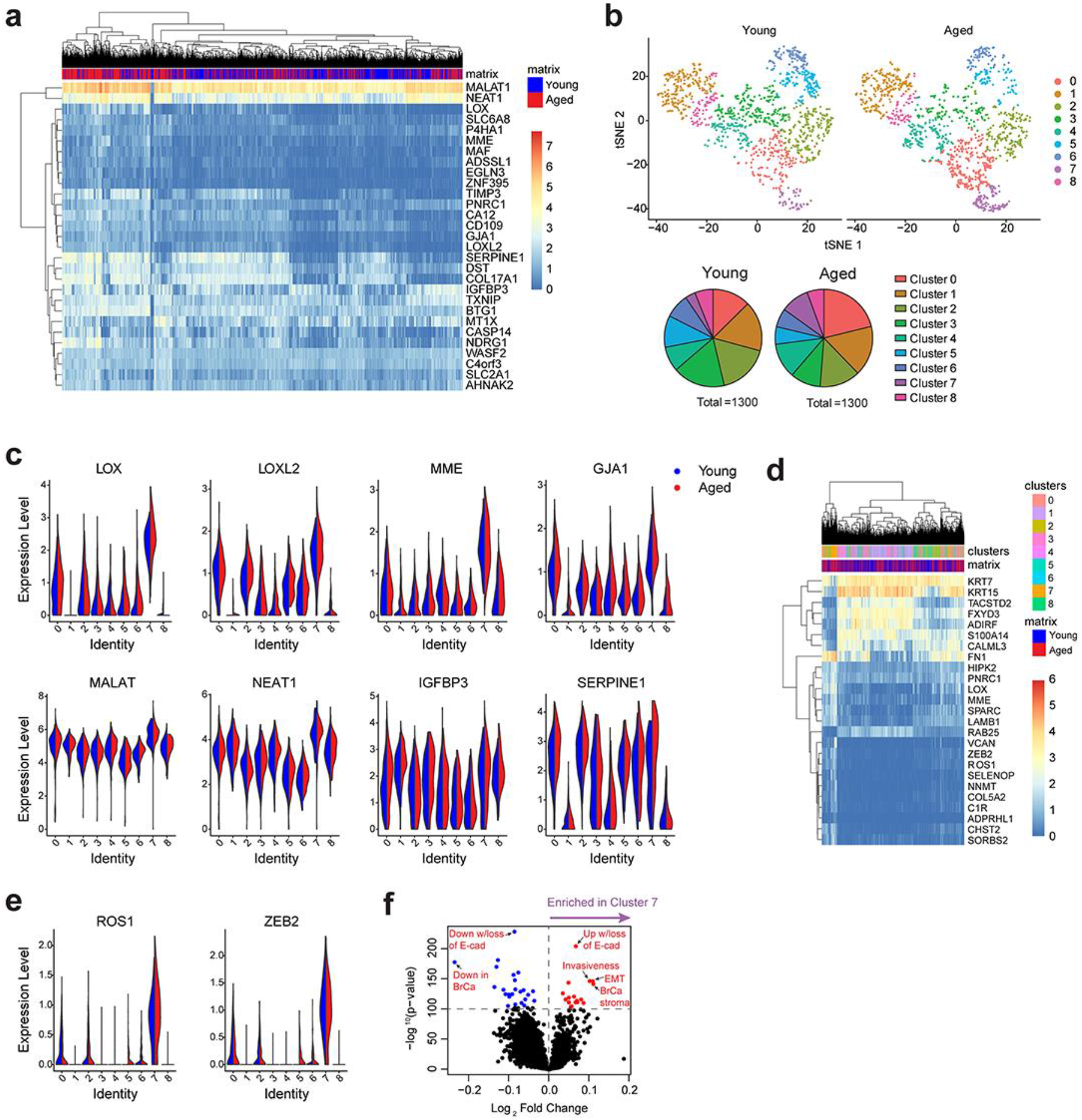
Single-cell RNA sequencing of KTB-21 cells incubated for 20 days on young and aged matrices reveals enrichment of an invasive sub-population on the aged matrices. (a) Genes significantly upregulated in cells on the aged matrix (adjusted p<0.05). (b) Clustering of cells on each particular matrix after down-sampling. Top: dimension reduction using t-distributed stochastic neighbor embedding (t-SNE) showing the distribution of cell clusters on the matrices. Bottom: pie chart showing the percentage of cells in each cluster. (c) Expression of the invasion and migration related genes that have upregulated on the aged matrices in cell clusters. (d) Genes significantly differentially expressed in cluster 7 cells compared to cells in other clusters. (e) Genes significantly upregulated in cluster 7 cells. (f) Single sample gene set enrichment analysis (ssGSEA) showing enrichment of invasion and EMT-associated gene sets in cluster 7 cells. n= 1300 cells/matrix type.

Closer examination of the genes that defined cluster 7 revealed that the cells were defined by loss of expression of the epithelial *TACSTD2* and *KRT15,* and the adipogenic *ADIRF* (Fig. 3d). More importantly, cluster 7 cells were defined by increased expression of a number of genes involved in EMT-like processes, including *LOX*, *ROS1*, *ZEB2*, *FN1*, *VCAN*, and *SPARC*. Specifically expression of *ROS1* and *ZEB2* was significantly higher in cluster 7 cells than others (Fig. 3e).

At the gene set signature level, single sample gene set enrichment analysis (ssGSEA) revealed enrichment of invasive and EMT-associated gene sets in cluster 7 cells (Fig. 3f). Taken together, single cell RNA-sequencing suggests the existence of a subpopulation of invasive cells in normal mammary epithelial cells, which is enriched when they engage with aged matrix compared to young.

### 2.4. LOX knockdown reverses the aged matrix-induced changes in cell phenotype, E-CAD localization, protein expression, and cell migration and invasion

LOX represses E-cadherin under hypoxic conditions^29^, and *LOX* knockdown stabilizes E-cadherin expression especially on the cell membrane^30^. As we showed that E-CAD was delocalized on cells in the aged microenvironment, and as *LOX* was highly expressed in cells on the aged matrices both in bulk and in cluster 7 cells, we investigated the effect of *LOX* siRNA treatment on E-CAD expression and localization, spheroid formation, cytokine production, and cell migration and invasion. *LOX* siRNA treatment resulted in reduced *LOX* mRNA (Figs. S5a and S5b) and protein (Fig. S5c) expression. Interestingly, E-CAD protein production was also reduced with *LOX* siRNA treatment. On the matrices, *LOX* siRNA treatment prevented the disappearance of KTB21 spheroids (red arrowheads) on the aged matrix until day 15 of culture, which otherwise disappear completely before day 15 (usually around day 13), and increased the number of spheroids on the young matrix (Fig. 4a). Scramble siRNA treatment did not show any difference from no treatment group (Fig. 4a vs Fig. 1b); spheroids on the aged matrix had disappeared by day 15, while those on the young matrix stayed intact. Additionally, *LOX* siRNA treatment reversed the delocalization of E-CAD on the aged matrix, leading to membrane-localized E-CAD expression, while slightly delocalizing E-CAD on the young matrix (Figs. 4a and 4b). LOX protein expression was reduced on the aged matrix after *LOX* siRNA treatment (p=0.0108) (Figs. 4a and 4c), showing successful knockdown of LOX in cells on these matrices. Interestingly, MMP2 (p=0.0073), but not MMP9, was significantly reduced on the aged matrix after *LOX* siRNA treatment.

**Fig. 4.**
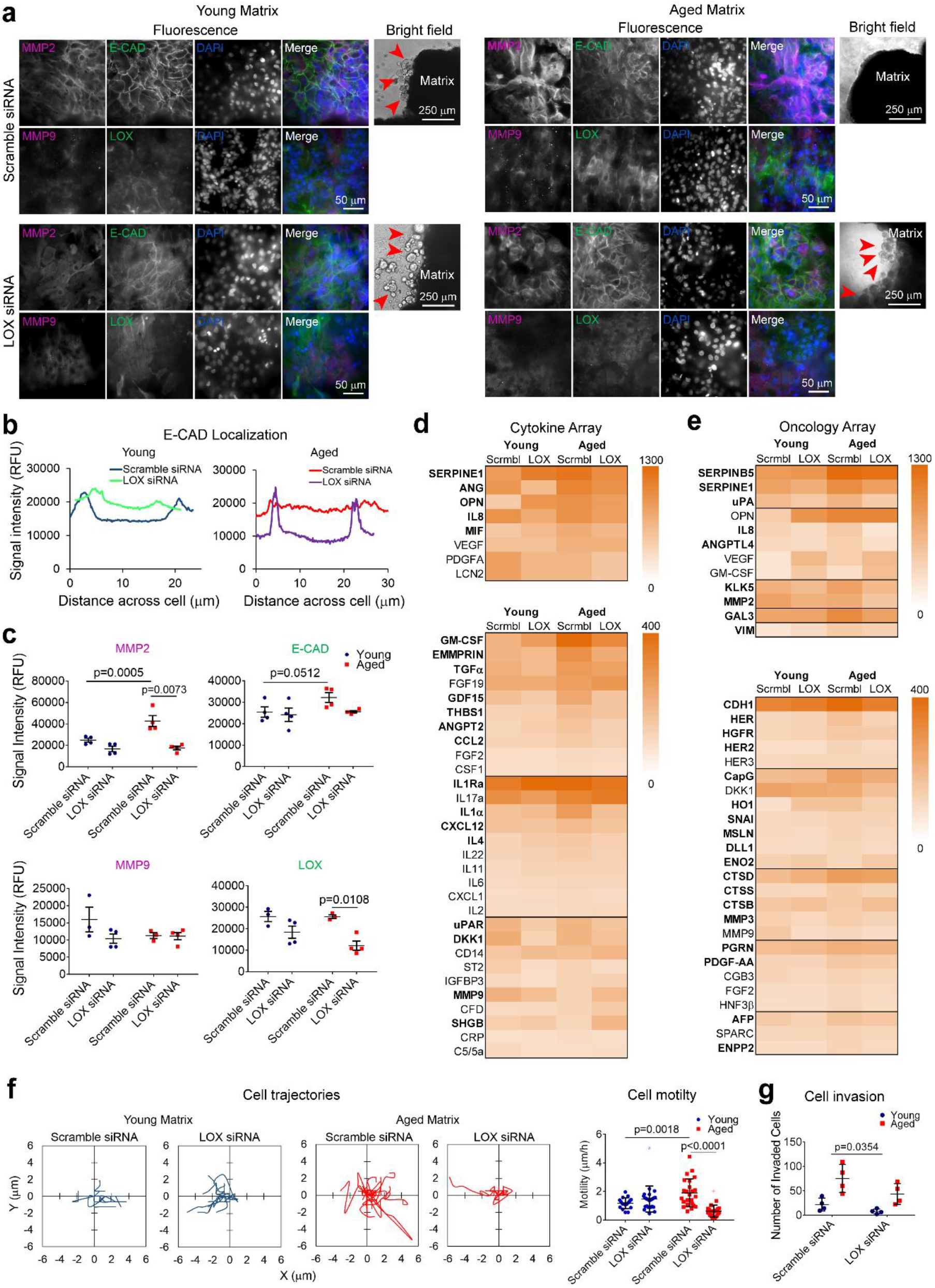
LOX knockdown reverses the phenotypic, biochemical and behavioral changes in KTB21 cells induced by the aged microenvironment. (a) MMP2, MMP9, E-CAD, and LOX expression, and spheroid stability on matrices (day 15). siRNAs applied between days 8-10. Fluorescence microscopy images showing MMP2 and MMP9 (magenta), E-CAD and LOX (green), and nucleic acid (DAPI, blue) staining, and merged images (Lane 4). Bright field images (Lane 5) showing the spheroids on matrices. Left: young matrices, and right: aged matrices. Upper panel: scramble siRNA, and lower panel: *LOX* siRNA. Arrowheads show spheroids. n=2 matrices. Results representative of two independent experiments. (b) Plot profile of E-CAD signal intensity along the diameter of representative cells in (a) showing E-CAD localization in the cells. n=4 images/matrix, and 8-10 cells/image. (c) Cell number-normalized signal intensities of MMP2, MMP9, E-CAD, and LOX as quantified from (a). n=2 matrices, 2 images/matrix. (d, e) Heat map showing the dot blot-based cytokine profiling of the cells on matrices (day 15). (d) Cytokine array, and (e) oncology array. Also see Tables S2 and S3. (f, g) Migration of KTB21 cells after *LOX* knockdown. siRNAs were applied on cells for 48 h before seeding on matrices. Time-lapse image taken at 15 min intervals for 2 h. (f) Cell trajectories (left) and the calculated motility results (right). (g) Invaded cell number after siRNA treatment. Cell-seeded matrices were placed in transwell inserts at day 7 and incubated against a 10% FBS gradient until day 21. siRNAs were applied between days 8-10, and also between days 15-17. Quantifications are performed using the Fiji software. Data are presented as the mean ± SD. Statistical tests: one-way ANOVA followed by Tukey’s post hoc for (c) and (f), and two-way ANOVA for (g).

We next performed cytokine profiling for the KTB21-seeded decell/delip matrices after siRNA treatment. Most of the cytokines, especially the uPA/uPAR mediators SERPINE1 (PAI-1), ANG, OPN, and uPAR, the pro-inflammatory IL8, MIF, GM-CSF, TNFα, IL1, and IL4, and the MMP enhancer EMMPRIN were produced at higher levels on the aged matrix, but were reduced to the young matrix levels after *LOX* siRNA treatment (Figs. 4d and S6a, and Table S2). Interestingly, some factors like FGF-19, IL17a, CD14, MMP9 and SHBG increased in the aged microenvironment after *LOX* siRNA treatment. We also performed oncology profiling to show how aging environment influenced the expression of some cancer-associated proteins. Similar to the cytokine profiling results, most of the cancer-associated proteins, such as ANGPTL4, KLK5, MMP2, GAL3, VIM, HERs, CAPG, HO, CTSD, and AFP, were produced at higher levels on the aged matrix, but were reduced to young matrix levels after *LOX* siRNA treatment (Figs. 4e and S6b, and Table S3). SERPINB5 (Maspin) and VEGF were produced at high levels on the aged matrix. Upon *LOX* siRNA treatment, VEGF was elevated, while SERPINB5 did not change.

Finally, we studied the effect of *LOX* knockdown on cell migration and invasion. While *LOX* knockdown showed no effect on the migration of KTB21 cells on the young matrix, it led to a significant reduction in cell migration on the aged matrix (Fig. 4f). Cell motility was even below the level of cells on the young matrix. Similar finding was observed with the cell invasion results. The number of invaded cells after 14-day incubation in the transwell inserts against 10% FBS gradient (following a 7-day pre-incubation in culture plates) decreased significantly upon *LOX* knockdown both on the young and on the aged matrices (p<0.0354, two-way ANOVA) (Figs. 4g and S7).

We also tested the effect of *LOX* siRNA on MDA-MB-231 cell migration and invasion behavior. Motility of the scramble siRNA-treated cells was significantly greater on the aged matrix than on the young (p=0.0018), and *LOX* siRNA treatment reduced the motility below the young control level (p<0.0001) (Figs. S8a and S8b). Cell invasion after 4-day incubation in transwell inserts against 10% FBS gradient (following a 7 day pre-incubation on culture plates) was also reduced on the young and aged matrices after *LOX* knockdown (p<0.0371, two-way ANOVA) (Fig. S8c).

## 3. Discussion

Here we report that aged mouse breast ECM promotes EMT-like and invasive behavior in normal (KTB21) and cancerous (MDA-MB-231) human mammary epithelial cells. Aged microenvironment induces E-CAD delocalization from cell membrane to cytosol, increases the expression of MMP2, pro-inflammatory cytokines (IL8, MIF, GM-CSF, TNFα, IL1, and IL4), uPA/uPAR system components (SERPINE1 (PAI-1), ANG, OPN, and uPAR), and several cancer-related proteins (ANGPTL4, KLK5, MMP2, GAL3, VIM, HERs, CapG, HO, CTSD, and AFP), promotes cell motility and invasion by upregulating *LOX*, *LOXL2*, *SERPINE1*, *MME*, *GJA1*, *MALAT*, *NEAT1*, and *IGFBP3*, and enriches a subpopulation of cells expressing EMT-related genes, including *ZEB2*, *ROS1*, *FN1*, *VCAN*, and *SPARC*. Remarkably, *LOX* knockdown leads to re-localization of E-CAD to cell membrane, represses the pro-inflammatory cytokines, oncogenic proteins, and the uPA system components, and reduces the cell motility on the aged matrix, matching the levels of cells in the aged microenvironment to those cells in the young.

Dissecting the reasons for the induced invasive and EMT-like phenotype of the normal and cancerous mammary epithelial cells in the aged microenvironment is difficult, since the ECM is a complex network of proteins and signaling molecules. We show that aging leads to thinner and more wavy collagen fibers, increased Young’s modulus, elevated levels of proinflammatory cytokines (Cd14, Cd40, Ccl21, Complement factor D, Il28, and Il33), protease inhibitors (Col18a1/Endostatin and Cystatin C), matrix metalloproteinases (MMP2 and MMP3), Igfbp3, and Igfbp6, and myeloperoxidase, as well as decreased levels of structural proteins (Col5 and Col15), each potentially contributing to EMT-like and invasive phenotype. Additionally, we show cytokines alone increase the invasiveness of cells, and along with other biochemical and physical cues, they may promote cell migration. Collagen structure (density and fiber thickness), stiffness, cytokines and chemokines, and biochemical composition may all play roles in cancer initiation and progression^1,11,17,23,24,31^. Here, we highlight the importance of decreased Col15, which is a tumor suppressor localized to basement membrane and involved in stabilization of E-CAD and prevention of its internalization to cytosol^15^. Loss of Col15 could be the reason for the delocalization of E-CAD in cells on the aged matrices. On the other hand, the reduced Col5 in the aged matrix, which plays a role in fibril assembly^32^, could be the reason for the thinner collagen fibers. A previous study has reported lower production of COL5 and COL15 proteins in xenografts created using the more aggressive MDA-MB-231-NM2 cell lines compared to the less aggressive MDA-MB-231^33^.

On the cellular side, we show that although KTB21 cells can form spheroids on both young and aged decell breast matrices, spheroids in the aged microenvironment deform and disappear between days 10-15 (usually around day 13) in culture. The disappearance of the spheroids in the aged microenvironment could be due to faster degradation of the matrix and/or weaker cell-cell adhesion. Indeed, we show that MMP2 is expressed at higher levels on the aged matrix and E-CAD is delocalized from cell membrane to cytosol. It is known that MMP2 expression is high in breast cancer^34,35^. Moreover, E-CAD internalization to cytosol plays role in EMT^36^, and its localization to cell membrane is disrupted in breast and other cancers^37,38^. Here, we show for the first time that aging microenvironment leads to E-CAD delocalization.

We next report increased motility and invasion of both normal and cancer cells, as well as upregulated *LOX*, *LOXL2*, *SERPINE1*, *MME*, *GJA1*, *MALAT1*, *NEAT1*, and *IGFBP3* expression by normal cells on the aged matrix, all associated with EMT, cell migration, and cancer invasion and progression^31,39,40^. Interestingly, a subset of cells (cluster 7) with highly upregulated EMT markers *ROS1*, *ZEB2*, *FN1*, *VCAN*, and *SPARC*, are also enriched in the aged breast microenvironment. Hence, here we show, for the first time, that the aged breast ECM alone can lead to enrichment of a subpopulation of normal epithelial cells with EMT-like and invasive phenotype.

LOX is an EMT marker^41^ reported to repress E-CAD^29^. However, here we rather show that LOX increases the expression of E-CAD in the aged matrix, as well as delocalizing it from cell membrane to cytosol. *LOX* mRNA is upregulated in cells on the aged matrices compared to young, and when it is knocked down, E-CAD in cells on the aged matrix is re-localized to cell membrane. Remarkably, cytokine levels in the aged microenvironment, especially the uPA/uPAR mediators uPA, uPAR, SERPINE1 (PAI-1), ANG, and OPN, which promote cell migration^42,43^, are reduced to the young matrix levels after *LOX* knockdown, while SERPINB5 (Maspin), which inhibits cell migration^44^, is not reduced. Loss of E-CAD from cell membrane is reported to increase the expression of uPAR^45^, supporting our finding. Our results also indicate that *LOX* knockdown leads to decrease in many factors including the intracellular (galectin 3, CapG, HO1, SNAI, mesothelin, and AFP), surface (E-cadherin, HER, HER2, and HGFR, thrombospondin 1) and extracellular (MMPs and cathepsins) proteins, cytokines (IL8, MIF, IL1α, IL4, CCL2), and growth factors (TGFα, GDF15, FGF, and angiopoietin 2), most of which are produced at higher levels in the aged microenvironment and involved in cell migration and invasion. However, *LOX* knockdown also leads to increased levels of the proangiogenic factors (VEGF, PDGF-AA, and GM-CSF), which may increase the risk of breast cancer progression.

As LOX is a collagen crosslinking enzyme and increases the stiffness of ECM, targeting LOX activity has previously been proposed to prevent metastasis^23,46^. LOX inhibitors such as β-aminopropionitrile (BAPN) and the aminomethylenepyridine based CCT365623 have proven effective in reducing metastasis in mouse models of breast^47,48^ and pancreatic^49^ cancers. However, it has been suggested that BAPN be administered before the tumor progresses to be effective, and otherwise would even lead to tumor progression^50^. As well as the active form of LOX enzyme, LOX inhibitors reduce the LOX propeptide, which plays role in inhibiting the pro-oncogenic β-catenin signaling through localizing β-catenin to cell membrane^51^. Therefore, targeting the active form of LOX instead of using non-specific anti-LOX drugs that also target the propeptide would be a more effective strategy in reducing breast cancer progression. Results of our study indicate potential benefits for the use of LOX inhibitors alone for the elderly as a preventative measure or with anti-angiogenic drugs for those who are at the early stages of cancer to prevent its invasion.

These findings indicate that the aged matrix create an invasive microenvironment for the cells. In line with our data, induced metastatic ability of the ovarian (OvCa)^52^ and prostate (TRAMP-C2 and Myc-CaP)^53^ cancer cells were reported in the aged microenvironment in mice. This study is important because it could lead to a better understanding of cell migration and invasion processes in aged tissues, which would lead to improved prognostic ability and disease outcome, as well as shedding light to cancer initiation and progression processes, since aged microenvironment seems to harbor components that could lead to cancer initiation or progression, while lacking other components that might prevent it. In the same vein, young microenvironment could shed light into new ways to prevent cancer. Analysis of how normal cell transcriptome changes in response to the aging ECM could enable identification of target genes that could be useful in preventing cancer initiation and progression. This study would usher the role of aging ECM not only in breast cancer, but also in many cancers and other age-related diseases, and pave the way for discovery of more efficient treatment strategies.

## 4. Materials and Methods

### 4.1. Breast tissue harvest, and decellularization/delipidation

Breast tissues (4^th^ mammary glands) were harvested from C57BL/6 mice at 3-6 (young) or 20-23 months of age (aged) according to the IACUC guidelines with the approval of the University of Notre Dame, which has an approved Assurance of Compliance on file with the National Institutes of Health, Office of Laboratory Animal Welfare. Mice were sacrificed in CO_2_ chambers, and tissues collected and used immediately, or wrapped in aluminum foil, flash frozen in liquid nitrogen, and stored at −80 °C until use.

To section the tissues in cryostat, tissues were thawed at room temperature, blotted on a tissue paper, and embedded in Tissue-Tek optimum cutting temperature (O.C.T.) compound (Sakura, USA) frozen at −20 °C, and sectioned at 300 μm thickness. Sections were placed in PBS to remove the O.C.T compound.

To remove cells (decellularization), whole tissues or sections were incubated in 0.5% SDS for 2-4 days at 4 °C, with gentle agitation and SDS change every 12 hours. The tissues were then briefly washed with PBS and incubated in 100% isopropanol for 2-4 days at 4 °C, with gentle agitation and isopropanol change every 12 hours to remove the lipids (delipidation). After decellularization/delipidation tissues were washed twice with PBS and stored at 4 °C until use (for tissue characterization matrices were used immediately, and for cell seeding experiments they were used within two weeks).

### 4.2. Mouse breast tissue characterization

#### 4.2.1. Microscopy

To analyze collagen structure, the native tissues, and decellularized (decell) and delipidized (delip) matrices were imaged using a two-photon confocal microscope (Olympus, FV1000 MPE) with second harmonic generation (SHG) at 800 nm wavelength, and the collagen fiber thickness was measured using ImageJ software (NIH, USA). n= 18 for the young and 21 for the aged native tissues, n=22 for the young and 21 for the aged decell matrices, and n=13 for the young and 15 for the aged delip matrices were used.

Surface properties of the native tissues were analyzed with digital filed emission scanning electron microscope (FEI Magellan 400, USA). Briefly, tissues sections were attached to stubs using carbon bands, coated with gold/palladium, and imaged under high vacuum, at 15 kV voltage.

#### 4.2.2. Nanoindentation testing

For the mechanical characterization, a nanoindenter (Piuma Chiaro, Optics11, The Netherlands) with a 10 N load cell, and a silicon nitride SNL-10 cantilever (Bruker, USA) with a spring constant of around 0.261 N/m was used. The native tissues, and the decell and decell/delip matrices (n=5 for young, and 5 for aged) were tested at 5-15 different locations with 5 measurements at each location. The loading velocity was 2 mm/s. Young's modulus was determined by a custom developed MATLAB code using Hertz contact model as described previously^19^, assuming a Poisson's ratio of 0.5.

#### 4.2.3. Cytokine profiling

To profile the cytokines in the mouse breast, whole native tissues or decell/delip matrices (n=3 pooled samples for each young and aged) were flash frozen in liquid nitrogen and ground using a mortar and pestle, and the resulting powders were suspended in protease inhibitor cocktail (Sigma, Cat No: P8340) solution and homogenized using an ultrasonicator. Next, Triton X-100 was added (final concentration: 1%) to disrupt cells and fatty components. The mixture was centrifuged 5 times at 10,000g for 5 minutes, with removal of the cellular debris and fatty components after each centrifugation cycle. Then, protein quantification was done using the bicinchoninic acid (BCA) rapid gold assay kit (Pierce, Thermo Fisher Scientific).

The relative content of 111 cytokine proteins (Table S1) was determined using the dot blot based mouse XL cytokine array kit following manufacturer’s instructions (R&D Systems, Cat. No: ARY028). Briefly, array membranes were blocked with an array buffer for 1h at RT, washed and incubated overnight at 4 °C in equal amounts of the tissue extracts. After washing step, the membranes were incubated in the biotinylated antibody cocktail solution for 1 h, in streptavidin-horseradish peroxidase (HRP) for 30 min, and in the chemiluminescent reagent for 1 min. Membranes were exposed to X-ray for 5-10 min using a biomolecular imager (ImageQuant LAS4000, GE Healthcare, USA). Relative cytokine content was determined after quantification of the dot intensity using ImageJ.

#### 4.2.4. Mass spectrometry

Dried breast tissues were lysed using a lysis buffer containing 6M urea 2M thiourea at 4 °C for 48-72 h, and ECM proteins were precipitated with cold acetone at −20 °C for overnight. The pellet was dissolved in 8M urea buffer containing protease inhibitors, followed by sonicating and centrifuging. 100 μg of protein was treated with 5 mM dithiothreitol for 25 min at 56 °C, and 14 mM iodoacetamide for 30 min at room temperature in dark. The protein mixture was then diluted with 25 mM Tris-HCl (pH 8.2) to achieve a final urea concentration of 1.8 M. Proteins were digested overnight with trypsin from bovine pancreas (Sigma-Aldrich, St Louis, MO, USA) at 37 °C at a 50:1 (m/m) protein to trypsin ratio in the presence of 1 mM CaCl_2_. The digestion reaction was stopped by adding trifluoroacetic acid to a final concentration of 0.4% (vol/vol). Samples were cleaned with ZipTip and re-suspended in MS loading buffer (1% HPLC grade acetonitrile (ACN), 0.1% formic acid (FA) in HPLC grade water).

The mass spectrometric analysis was performed on a Q-Exactive mass spectrometer (Thermo Fisher Scientific) coupled with a nanoACQUITY Ultra Performance LC (UPLC) system (Waters Corporation). Peptides were loaded onto a C18 reverse phase column (100μm×100mm, 1.7μm particle size, BEH130) (Waters Corporation) with 97% buffer A (0.1% FA in water) and 3% buffer B (0.1% FA in ACN). Peptide separation was carried out with a 73-min linear gradient from 3% to 40% buffer B. All samples were run in technical triplicates.

All .raw files acquired with the Q-Exactive were searched with the Global Proteome Machine (GPM). The peptide false discovery rate (FDR) was determined by searching against the corresponding reverse database. The searches were performed with precursor peptide mass tolerance of 10ppm and fragment ion mass tolerance of 0.02 Da, and allow up to two missed cleavages with trypsin digestion. Carbamidomethylation of cysteine was set as a fixed modification, while oxidation of methionine was set as a variable modification (FDR = 0.01).

#### 4.2.5. Immunohistochemistry and histology

Native tissues were fixed in paraformaldehyde solution (4%), embedded in paraffin, sectioned at 6 μm thickness, attached to positively charged Superfrost^TM^ plus gold slides (Thermo Fisher Scientific), and deparaffinized and rehydrated.

For collagen I staining, tissue samples were incubated for 5 min in 0.3% Triton X-100, for 45 min in 5% goat serum, overnight at 4 °C in rabbit anti-collagen I antibody (Abcam, USA, Cat. No: ab34710, dilution: 1:100.), and for 1 h in goat anti-rabbit IgG secondary antibody (Abcam, Cat. No: ab150077). The slide was covered with Prolong^®^ Gold antifade reagent with DAPI (Cell Signaling Technology, USA, Cat, No: 8961) prior to covering with a coverslip. Tissues were then imaged under an inverted fluorescence microscope (Zeiss Axio Observer.Z1).

For hematoxylin and eosin staining, slides were stained with Mayer’s hematoxylin for 2 min, washed in 1% acid alcohol (70% ethanol containing 1% hydrochloric acid) for 30 s, blued with ammonia water (0.2%), stained in eosin-phloxine solution (1%) for 10 min, dehydrated and mounted.

For Masson’s trichrome staining, slides were stained in Mayer’s hematoxylin solution for 2 min, in Biebrich scarlet-acid fuchsin solution for 10 min, in phosphomolybdic-phosphotungstic acid solution for 10 min, in aniline blue solution for 5 min, and in 1% acetic acid solution for 2-5 minutes. Tissues were imaged under a bright filed microscope (Nikon, Eclipse ME600, USA).

### 4.3. Cell seeding and culture

KTB21 human mammary basal epithelial cell line (immortalized by human telomerase gene (hTERT) retrovirus using the vector pLXSN-hTERT) was previously established in Dr. Harikrishna Nakshatri’s lab from a 40 year-old patient^28^. KTB21 cells were cultured in epithelial cell growth medium (DMEM (low glucose):Ham’s F12 (1:3) medium supplemented with 5% FBS (Thermo Fisher Scientific, USA), 0.4μL/mL hydrocortisone (Sigma, USA), 1% penicillin/streptomycin (Corning, USA), 5μg/mL insulin (Sigma), 10 ng/mL EGF (Millipore, USA), 6mg/mL Adenine (Sigma), and 10mM ROCK inhibitor (Y-27632) (Enzo Life Sciences, USA)). Before cell seeding decell/delip matrices were sterilized in a solution containing 4% ethanol and 0.15% peracetic acid, washed with PBS and then with media, and placed in 96-well culture plates. To induce KTB21spheroid formation matrices were coated with Matrigel, since no spheroid formation was observed on the Matrigel-free matrices (Fig. S9). Briefly, the matrices were coated with 20 μL Matrigel (growth factor reduced, phenol red-free, and LDEV-free) (Corning, USA) and incubated for 10 min at 37 °C, before cells were seeded. The cells were then incubated in epithelial cell growth medium, with media change every 2-3 days. Spheroid formation and stability were monitored for 15 days.

For cell migration and invasion experiments, Matrigel-free matrices were used. KTB21 and the GFP reporting MDA-MB-231.BR cells (a gift from Dr. Patricia Steeg at NCI) were seeded on the matrices, and incubated in epithelial and cancer cell growth medium (DMEM (high glucose) supplemented with 10% FBS and 1% penicillin/streptomycin), respectively, with media change every 2-3 days.

### 4.4. LOX siRNA treatment and confirmation of the knockdown

For knockdown experiments, scramble (Cy3-labeled negative control) (Thermo Fisher Scientific, Cat No: AM4621), and *LOX* (Thermo Fisher Scientific, Cat. No: 4390824) siRNAs were applied for 48 h. Briefly, siRNAs (80 nM in serum free medium) and the Lipofectamine RNAiMax transfection reagent (Thermo Fisher Scientific, Cat. No: 13778075) (12 μg/mL in serum free medium) were mixed at 1:1 volume ratios (final siRNA concentrations: 40 nM) and incubated for 5 min at room temperature to allow complex formation. The siRNA/Lipofectamine complex was then applied on cells on tissue culture polystyrene (TCPS) or on matrices after washing the cells with PBS. To verify that we knocked down the *LOX* mRNA and LOX protein, we performed quantitative reverse transcriptase-polymerase chain reaction (qRT-PCR) for KTB21 and MDA-MB-231 cells, and western blotting for KTB21 cells cultured on TCPS, respectively.

For qRT-PCR, RNAs were collected using the RNA isolation kit (RNeasy, Qiagen), and cDNAs were synthesized using the iScript cDNA Synthesis Kit (Bio-Rad). Quantitative PCR was done using the iTaq SYBR Green Supermix (Bio-Rad) with certified human GAPDH and LOX primers (Bio-Rad). The reaction was run in CFX Connect 96 Real Time PCR system (Bio-Rad). The ΔΔCq method was applied to quantify the relative expression of genes, and GAPDH was used as the housekeeping gene.

For western blotting, cells seeded in TCPS were incubated for 48 h in the siRNA solutions, followed by another 48 h incubation in culture media, cells were incubated in RIPA buffer for 30 min in ice to collect the proteins. Protein quantification was done using the BCA assay. The same amounts of proteins were loaded into polyacrylamide gels (8%) and the samples run at 125 V for 2 h. The proteins were transferred to nitrocellulose membranes and incubated overnight in mouse anti-LOX (LSBio, Cat. No: LS-C172331), mouse anti-E-CAD (Abcam, Cat. No: ab1416), and rabbit anti-beta actin (Cell Signaling Technology, Cat. No: 3700) at 1:500 dilutions, followed by a 1 h incubation in the HRP-labelled secondary antibodies (horse anti-mouse IgG (Cell Signaling Technology, Cat. No: 7076), and goat anti-rabbit IgG (Cell Signaling Technology, Cat. No: 7074) at 1:1000 dilutions. Next, the membranes were incubated in chemiluminescent substrate (SuperSignal West Pico PLUS, Thermo Fisher Scientific) for 5 min and imaged.

For KTB21 cell spheroid experiments, siRNAs were applied between days 8 and 10 after cell seeding on matrices. For cell invasion experiments, siRNAs were applied between days 1 and 3, and day 8 and 10 after cells were seeded on the matrices, just before invasion assays were started. For all cell migration experiments, siRNAs were applied on TCPS, just before cell seeding on matrices.

### 4.5. Spheroid formation and protein deposition

Viability of KTB21 cells on the matrices was determined using the Live/dead cell viability assay kit (Thermo Fisher Scientific). Briefly, cells were incubated in a solution of calcein-AM (2 μM) (live cells, green) and of ethidium homodimer-1 (EthD-1) (4 μM) (dead cells, red) at 37 °C for 30 minutes. After incubation, cells were visualized using the fluorescence microscope. Cell viability was calculated using ImageJ software based on the living cell number divided by the total cell number.

Spheroid morphology was analyzed from the live/dead images using the shape descriptor analysis in ImageJ, and circularity and aspect ratio were estimated.

After 15-day culture of KTB21 cells on the matrices, samples were analyzed with a multiphoton microscope at 800 nm wavelength (second harmonic generation, SHG) to visualize the collagen in spheroids. Untreated (no siRNA) samples were stained for E-CAD or double stained for MMP2 and COL1. *LOX* siRNA-treated samples were double stained for E-CAD and MMP2 or for LOX and MMP9.

Briefly, whole samples were incubated for 15 min in 4% paraformaldehyde solution, for 5 min in 0.3% Triton X-100, and for 45 min in 5% goat serum, with three washes in PBS each for 5 min between every step. The samples were incubated overnight in mouse anti-E-CAD (Abcam, Cat. No: ab1416), mouse anti-COL1 (Abcam, Cat. No: ab90395), rabbit anti-MMP2 (CusaBio, Cat. No: CSB-PA06879A0Rb), rabbit anti-MMP9 (Abcam, Cat. No: ab76003), and mouse anti-LOX (LSBio, Cat. No: LS-C172331) with 1:100 dilutions, and then in Alexa fluor 488-labelled goat anti-mouse IgG (Abcam, Cat. No: ab150117, dilution: 1:400), and Alexa fluor 647-labelled goat anti-rabbit IgG (Abcam, Cat. No: ab150083, dilution: 1:400) secondary antibodies. The samples were stained with 1 mg/mL DAPI (Sigma, Cat, No: 10236276001) and imaged with a multiphoton microscope or an inverted fluorescence microscope.

### 4.6. Migration and invasion assays

For migration assays, untreated, scramble siRNA and *LOX* siRNA treated KTB21 and MDA-MB-231 cells were used. For siRNA knockdown experiments, cells were seeded on TCPS, grown until around 70% confluence, and treated with siRNA for 48 h. To track KTB21 cells under microscope, they were washed with PBS, stained with 10 μm CellTracker Green 5-chloromethylfluorescein diacetate (CMFDA) (Thermo Fisher Scientific) for 30 min. MDA-MB-231 cells are already GFP reporting.

Cells were seeded on the matrices at 2 × 10^6^ cells/mL (around 3 × 10^4^ cells/matrix) density. After 24 h, cells were imaged with fluorescence microscope at 15 min intervals for 4-20 h. Time=lapse images were concerted to .AVI video files and analyzed using the MTrack plugin in ImageJ to track the cells. Cell trajectories and motility (migration speed), as well as cell morphology (circularity and roundness) were calculated based on the distance travelled by the cells and the time during which cells were tracked.

Similarly, for invasion assays, untreated, scramble siRNA and *LOX* siRNA treated KTB21 and MDA-MB-231 cells were used. Briefly, cells were seeded on matrices at 4 × 10^6^ cells/mL (around 5 × 10^4^ cells/matrix) density, and cultured for 5 days until they reached confluence. At day 5 of culture, siRNAs were applied (for the knockdown groups), and the matrices were transferred at day 7 to transwell inserts (with 8 μm pores) (Corning, Cat. No: 29442-120) at day 7. Invasion assays were started at day 7, and terminated at day 14 (no treatment group) or 21 (siRNA treatment groups) for KTB21 cells, and at day 11 for MDA-MB-231 cells. In the transwell (upper) chamber, KTB21 cell-seeded matrices were incubated in regular epithelial growth medium (5% FBS), while MDA-MB-231 cell-seeded matrices were incubated in serum-free DMEM. A 10% FBS gradient was applied; in the bottom chamber epithelial growth medium with 15% FBS was used for KTB21 cells, while cancer cell growth medium (10% FBS) was used for MDA-MB-231 cells.

Alternatively, to test the effect of cytokines in the matrices KTB21 cells (10^4^ cells/well) were seeded in the transwell and the young and aged matrices were placed in the bottom chambers as incentives for invasion. Cells were incubated for 14 days. At the end of invasion tests, cells that invaded to bottom chambers were counted.

### 4.7. Single-cell RNA-sequencing

KTB21 cells were incubated on the Matrigel-coated matrices for 20 days. Three samples for each matrix type were pooled and treated with trypsin (0.25%) to collect the cells. Preparation for scRNA-seq was performed as described elsewhere^54^. Briefly, cells were resuspended in calcium-and magnesium-free PBS containing 2% BSA and 0.02% Tween 20 at 1 million cells/mL density, and blocked by adding 10 μL Human TruStain FcX blocking solution (Biolegend, 422301) and incubating on ice for 20 min. Next, 0.3 μL human hashtag (HTO) antibody (Biolegend) was added to each sample (a different hashtag antibody was used for each sample/replicate) and incubated on ice for 25 min. Finally, cells were washed four times in a series of buffers (first wash: 2% BSA, 0.02% Tween 20 in PBS; second wash: 2 mM EDTA, 2% BSA, 0.02% Tween 20 in PBS; third wash: 1 mM EDTA, 2% BSA, 0.02% Tween 20 in PBS; fourth wash: 0.1 mM EDTA, 1% BSA, and 0.02% Tween 20 in PBS). Samples were pooled prior to the final wash and counted, and following the final wash, resuspended at 1500 cells/μL.

10x Genomics Chromium was used for single cell capture, cDNA libraries were prepared according to the standard CITE-seq and 10x Genomics standard protocols. The resulting HTO-derived and mRNA-derived cDNA libraries were pooled and sequenced. 26 bp of cell barcode and UMI sequences and 91 bp RNA reads were generated with Illumina NovaSeq 6000. The raw base sequence calls generated from the sequencer were demultiplexed into sample-specific mRNA, ADT and HTO FASTQ files with bcl2fastq through CellRanger 3.1.0. Raw FASTQ files were processed using Cellranger 3.1.0.

Data analysis was performed in R (v 3.6.2) using the Seurat package (v 3.1.2)^55,56^ for data normalization, dimension reduction, clustering, and differential gene expression analysis. Samples were demultiplexed by HTO expression with a positive quantile of > 0.99. For quality control, cells with greater than 1,000 mRNA transcripts, and less than 20% mitochondrial genes were kept for analysis. To ensure even comparisons between cells cultured on young and aged matrices, each sample was randomly subsetted to include 1,300 cells. Dimension reduction and clustering was performed using standard parameters for the combined young and aged matrix datasets, followed by determination of the number of cells from each experimental condition present in each cluster. ssGSEA analysis was performed using GSVA (v 1.34.0) with all C2 gene sets from the Molecular Signatures Database (MSigDB).

### 4.8. Cytokine and oncology arrays

For cytokine analysis, the dot blot based human XL cytokine array kit (R&D Systems, Proteome profiler, Cat. No: ARY022B), which detects 105 cytokine proteins (Table S2), and human XL oncology array kit (R&D Systems, Proteome profiler, Cat. No: ARY026), which detects 84 human cancer-related proteins (Table S3), were used as described above (section 4.2.3). Supernatants of the KTB21 cell-seeded, Matrigel-coated matrices were collected at day 15, and the protein quantification was done using the BCA rapid gold assay kit. The same amounts of proteins were loaded onto the membranes and relative cytokine content was determined after quantification of the dot intensity using ImageJ.

### 4.9. Statistical analysis

Data were analyzed for statistical significance using GraphPad Prism 6 or R software. One-way analysis of variance (ANOVA) followed by Tukey’s HSD or two-tailed Student’s t-test were performed to compare the results. Two-way ANOVA was applied to test the effect of two independent variables (age and siRNA treatment) on a dependent variable (motility and invading cell number), and whether there is an interaction between the two independent variables. Data are presented as the mean ± standard deviation (SD).

## Acknowledgements

This study is funded by NIH award number 5R01EB027660-02 and Walther Cancer Foundation, Harper Cancer Research Institute Cancer Cure Ventures Award number 0184.01. EH is supported by NIH through grant number F32 CA210583-01. MSS is supported by NIH through UO1 CA236979.

## Author contribution

GB, XY, SZ and PZ designed the study. GB wrote the manuscript. GB, XY, and IG performed the experiments. GB, XY, and EH analyzed the data. SZ, HN, SS and PZ provided resources. SZ and PZ provided funding. All authors reviewed and revised the manuscript.

