## Supplementary Files for "Aged Breast Extracellular Matrix Drives Mammary Epithelial Cells to an Invasive and Cancer-Like Phenotype"

### Supplementary Materials

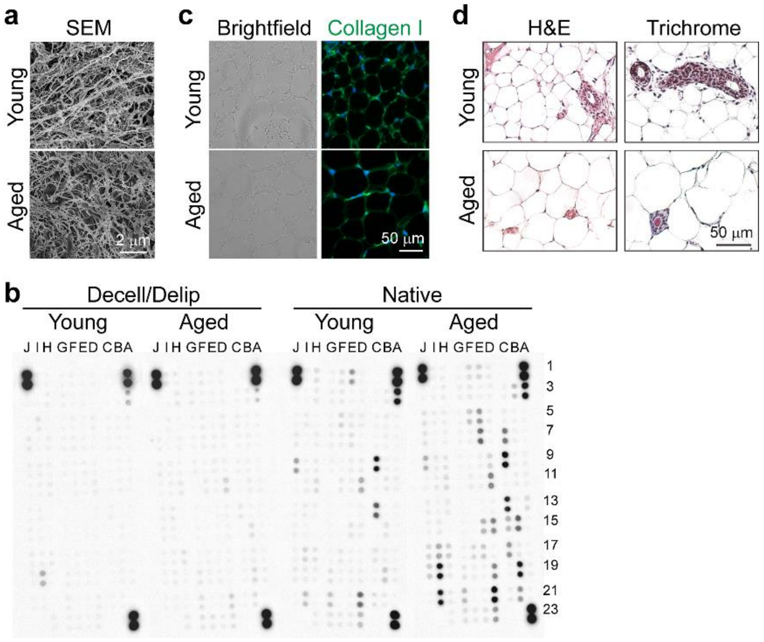


**Fig. S1.** Aging microenvironment alters the structure, biochemistry, and histology of the breast tissue. (a) Scanning electron microscopy images of the native breast tissues. (b) Dot blot of mouse cytokines. Also see Table S1. A1 and A2, A23 and A24, and J1 and J2 are reference spots, and J23, and J24 are negative controls. (c) Bright field (left) and fluorescence (right) microscopy images after staining for collagen. Green: collagen 1 (Alexa fluor 488), and blue: nuclei (DAPI). (d) Bright field microscopy images showing histology staining. Left: hematoxylin and eosin (H&E) staining, and right: trichrome staining.

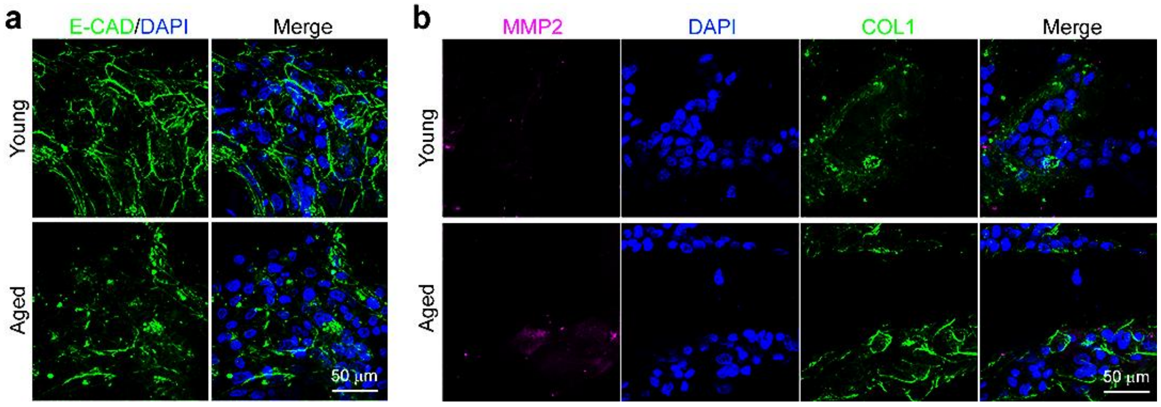


**Fig. S2.** E-CAD, MMP2 and COL1 expression in KTB21 cells. (a) E-CAD expression showing the localization cells. Green: E-cadherin (Alexa fluor 488). (b) MMP2 and COL1. Magenta: MMP2, green: COL1. Blue: nuclei (DAPI).

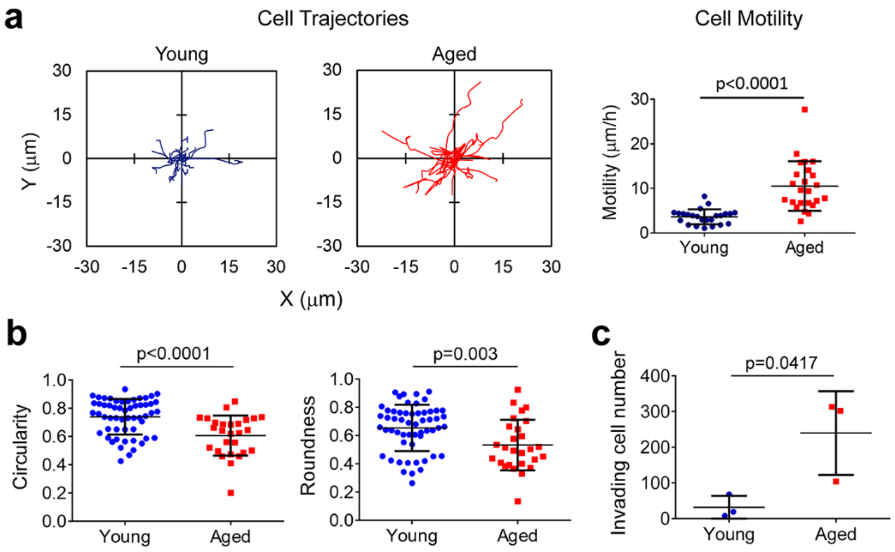


**Fig. S3.** Aged microenvironment enhances the motility and invasiveness of the MDA-MB-231 breast cancer epithelial cells, and changes their morphology. (a, b) Migration behavior of the MDA-MB-231 cells on the Matrigel-free matrices. Time-lapse images of the cells were taken at 15 min intervals for 3 h after incubating the cells for one day on the matrices. (a) Cell trajectories (left), and motility (left). (b) Morphology of the cells during migration on the matrices. Left: circularity, and right: roundness. (c) Cell invasion through transwell inserts. Cells were seeded on the matrices and pre-incubated for 7 days, and then the matrices were incubated in transwell inserts for 4 days against a 10% FBS gradient. n=3-4 matrices.
Quantifications were performed using the Fiji software. Data are presented as the mean ± standard deviation. Statistical tests: student's t-test.

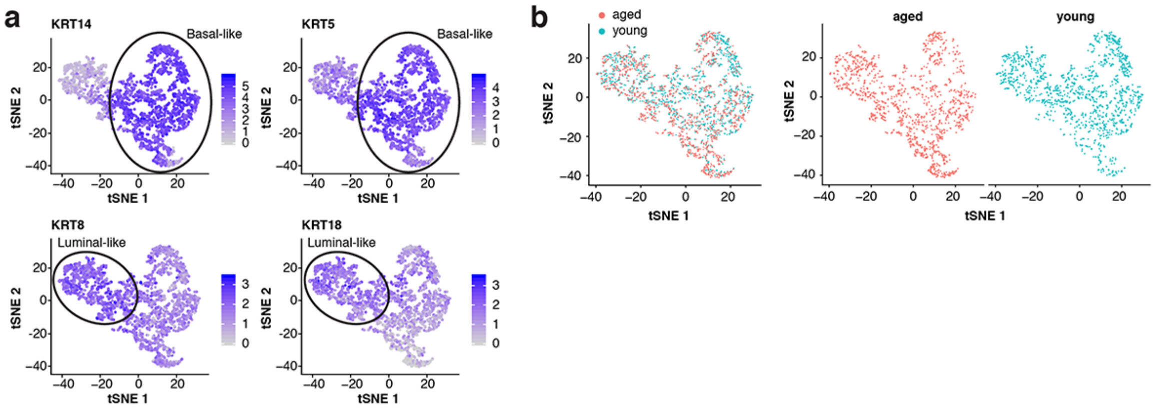


**Fig. S4.** KTB21 basal epithelial cells maintain their basal phenotype. (a) t-SNE showing the expression of basal epithelial markers, KRT5 and KRT14, and luminal epithelial markers, KRT8 and KRT18. (b) t-SNE showing differential gene expression of all cells on the young and aged matrices.

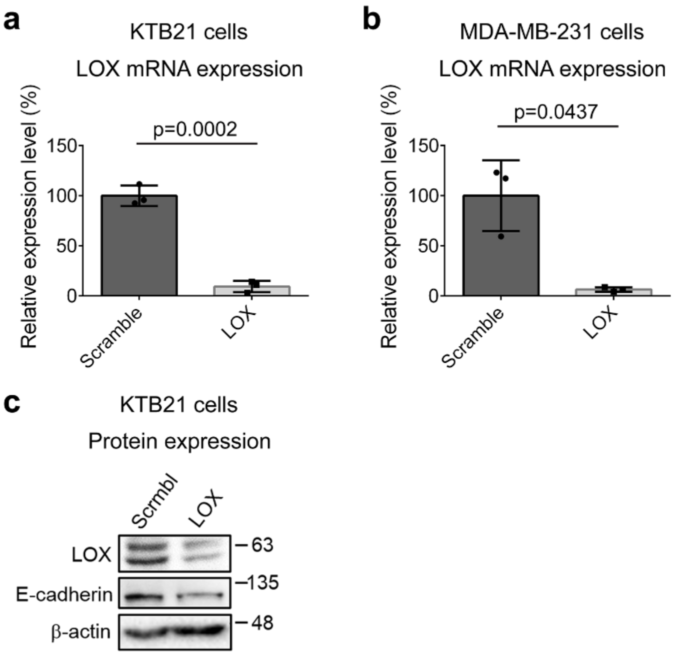


**Fig. S5.** mRNA and protein expression after *LOX* siRNA treatment. (a,b) qRT-PCR results showing the *LOX* mRNA expression after scramble and *LOX* siRNA treatment. (a) KTB21 cells, and (b) MDA-MB-231 cells. (c) Western blots showing the LOX and E-cadherin protein expression after scramble and *LOX* siRNA treatment. β-actin was used as the reference protein. Student's t tests were applied to test the difference between scramble and *LOX* siRNA treatment groups.

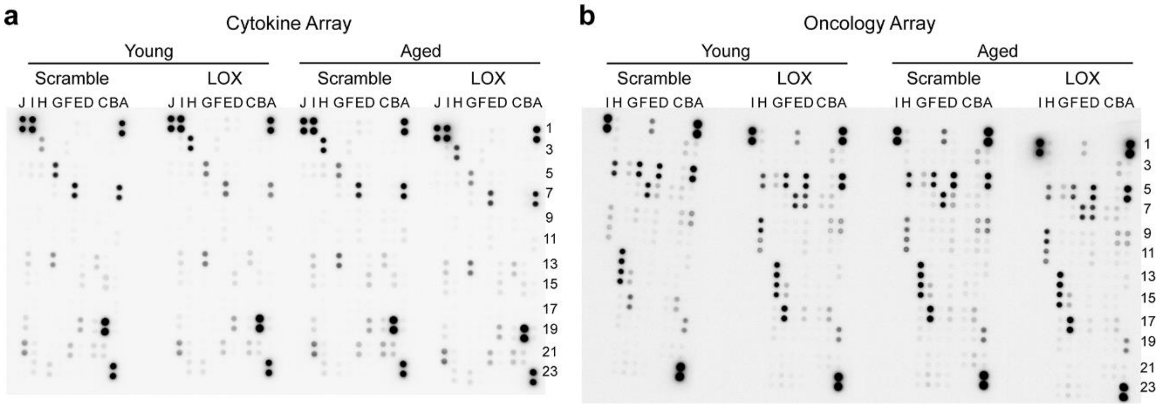


**Fig. S6.** Dot blots of the KTB21 cell lysates after 15 day incubation on the decell/delip matrices showing the effect of LOX knockdown on cytokine production. (a) Cytokine array, and (b) oncology array results of the KTB21 cells cultured for 15 days on matrices. siRNAs were applied between days 8-10 for 48 h. For (a) also see Table S2. A1 and A2, A23 and A24, and J1 and J2 are reference spots, and J23, and J24 are negative controls. For (b) also see Table S3. A1 and A2, A23 and A24, and I1 and I2 are reference spots, and I23, and I24 are negative controls.

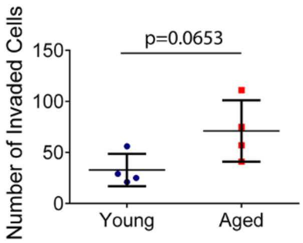


**Fig. S7.** Invasion of KTB21 cells on young and aged matrices. Invaded cell numbers. Cell-seeded matrices were placed in transwell inserts at 7 days after seeding and incubated against a 10% FBS gradient until day 21.

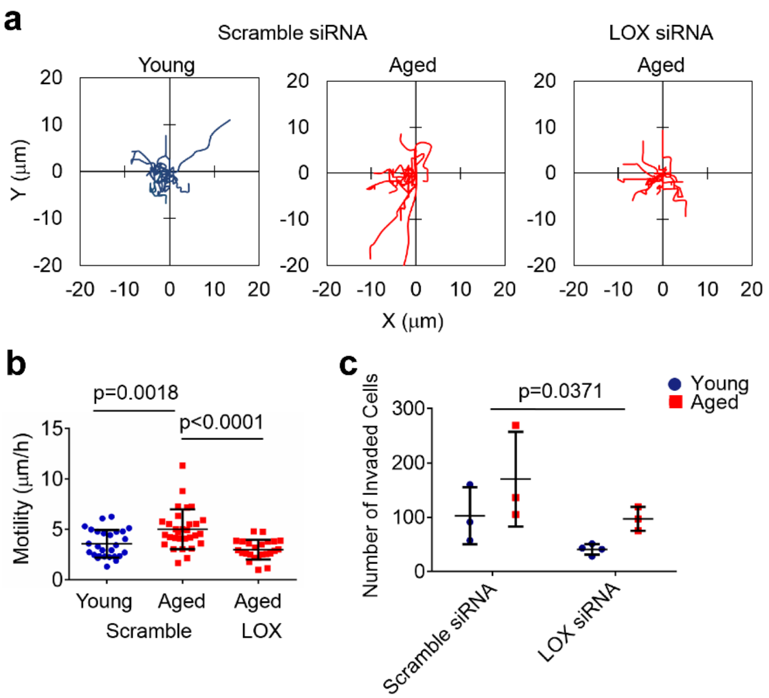


**Fig. S8.** Knockdown of LOX gene reduces the motility and invasiveness of MDA-MB-231 cells on the aged matrices to the level of young matrices. (a, b) Migration behavior of the cells after 1 day incubation on the matrices. (a) Cell trajectories, and (b) motility. Cells were incubated in the siRNAs for 48 h, and then seeded on the matrices. After 1 day incubation on the matrices, time-lapse images of the cells were taken at 15 min intervals for 3 h. (c) Invasion of the cells after scramble and *LOX* siRNA treatment. Cells were seeded on the matrices, incubated for 5 days, treated with siRNAs for 48 h between days 5 and 7. Invasion assay was started at day 7 of culture and applied for 4 days. n=3-4 matrices.

Quantifications were performed using the Fiji software. Data are presented as the mean ± standard deviation. Statistical tests: (b) one-way ANOVA followed by Tukey's post hoc; (c) two-way ANOVA.

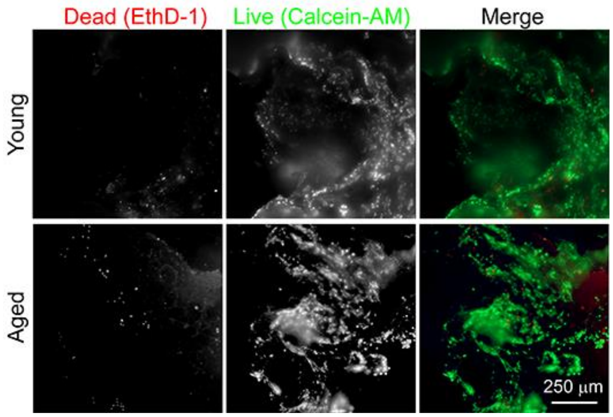


**Fig. S9.** KTB21 cells cannot form spheroids on the decellularized matrices. Fluorescence microscopy images of live/dead stained KTB21 cells at day 14 of incubation on the decell/delip matrices. Red: dead cells (ethidium homodimer-1 (EthD-1)), and green: live cells (calcein-AM).

**Table S1.** Mouse cytokines tested in dot blot assay.

| Spot Location | Protein | Spot Location | Protein | Spot Location | Protein |
|---|---|---|---|---|---|
| **A1, A2** | Reference Spots | **D15, D16** | Cd26 | **G21, G22** | Il-33 |
| **A3, A4** | Adiponectin | **D17, D18** | Egf | **G23, G24** | Ldlr |
| **A5, A6** | Amphiregulin | **D19, D20** | Endoglin | **H1, H2** | Leptin |
| **A7, A8** | Angiopoietin-1 | **D21, D22** | Endostatin | **H3, H4** | Lif |
| **A9, A10** | Angiopoietin-2 | **D23, D24** | Fetuin A (Ahsg) | **H5, H6** | Lipocalin-2 |
| **A11, A12** | Angiopoietin-like 3 | **E1, E2** | Fgf-a | **H7, H8** | Lix |
| **A13, A14** | Baff | **E3, E4** | Fgf21 | **H9, H10** | M-Csf |
| **A15, A16** | Cd93 | **E5, E6** | Flt-3 ligand | **H11, H12** | Mmp2 |

| | | | | | |
|---|---|---|---|---|---|
| **A17, A18** | Ccl2 | **E7, E8** | Gas 6 | **H13, H14** | Mmp3 |
| **A19, A20** | Ccl3 | **E9, E10** | G-CSF | **H15, H16** | Mmp9 |
| **A21, A22** | Ccl5 | **E11, E12** | Gdf15 | **H17, H18** | Myeloperoxidase (Mpo) |
| **A23, A24** | Reference Spots | **E13, E14** | Gm-Csf | **H19, H20** | Osteopontin (Opn) |
| **B3, B4** | Ccl6 | **E15, E16** | Hgf | **H21, H22** | Osteoprotegerin |
| **B5, B6** | Ccl11 | **E17, E18** | Icam-1 | **H23, H24** | Pd-Ecgf |
| **B7, B8** | Ccl12 | **E19, E20** | Ifn-$\gamma$ | **I1, I2** | Pdgf-bb |
| **B9, B10** | Ccl17 | **E21, E22** | Igfbp1 | **I3, I4** | Pentraxin 2 |
| **B11, B12** | Ccl19 | **E23, E24** | Igfbp2 | **I5, I6** | Pentraxin 3 |
| **B13, B14** | Ccl20 | **F1, F2** | Igfbp3 | **I7, I8** | Periostin |
| **B15, B16** | Ccl21 | **F3, F4** | Igfbp5 | **I9, I10** | Pref1 |
| **B17, B18** | Ccl22 | **F5, F6** | Igfbp6 | **I11, I12** | Proligerin |
| **B19, B20** | Cd14 | **F7, F8** | Il-1$\alpha$ | **I13, I14** | Pcsk9 |
| **B21, B22** | Cd40 | **F9, F10** | Il-1$\beta$ | **I15, I16** | Rage |
| **C3, C4** | Cd160 | **F11, F12** | Il-1ra | **I17, I18** | Rbp4 |
| **C5, C6** | Chemerin | **F13, F14** | Il-2 | **I19, I20** | Reg3G |
| **C7, C8** | Chitinase 3-like 1 | **F15, F16** | Il-3 | **I21, I22** | Resistin |
| **C9, C10** | Tissue factor | **F17, F18** | Il-4 | **I23, I24** | Reference Spots |
| **C11, C12** | Complement component C5/C5a | **F19, F20** | Il-5 | **J1, J2** | Reference Spots |
| **C13, C14** | Complement factor D | **F21, F22** | Il-6 | **J3, J4** | E-Selectin |

| | | | | | |
|---|---|---|---|---|---|
| **C15, C16** | C-reactive protein | **F23, F24** | Il-7 | **J5, J6** | P-Selectin |
| **C17, C18** | Cx3cl1 | **G1, G2** | Il-10 | **J7, J8** | SerpinE1 (PAI-1) |
| **C19, C20** | Cxcl1 | **G3, G4** | Il-11 | **J9, J10** | SerpinF1 |
| **C21, C22** | Cxcl2 | **G5, G6** | Il-12 p40 | **J11, J12** | Thrombopoietin |
| **D1, D2** | Cxcl9 | **G7, G8** | Il-13 | **J13, J14** | Tim-1 |
| **D3, D4** | Cxcl10 | **G9, G10** | Il-15 | **J15, J16** | Tnf$\alpha$ |
| **D5, D6** | Cxcl11 | **G11, G12** | Il-17A | **J17, J18** | Vcam-1 |
| **D7, D8** | Cxcl13 | **G13, G14** | Il-22 | **J19, J20** | Vegf |
| **D9, D10** | Cxcl16 | **G15, G16** | Il-23 | **J21, J22** | Wisp-1 |
| **D11, D12** | Cystatin C | **G17, G18** | Il-27 p28 | **J23, J24** | Negative Control |
| **D13, D14** | Dkk-1 | **G19, G20** | Il-28A/B | | |

**Table S2.** Human cytokines tested in dot blot assay.

| **Spot Location** | **Protein** | **Spot Location** | **Protein** | **Spot Location** | **Protein** |
|---|---|---|---|---|---|
| **A1, A2** | Reference Spots | **D11, D12** | IGFBP2 | **G13, G14** | MIF |
| **A3, A4** | Adiponectin | **D13, D14** | IGFBP3 | **G15, G16** | MIG |
| **A5, A6** | Apolipoprotein A-I | **D15, D16** | IL1$\alpha$ | **G17, G18** | MIP1$\alpha$/MIP1$\beta$ |
| **A7, A8** | Angiogenin | **D17, D18** | IL1$\beta$ | **G19, G20** | MIP3$\alpha$ |
| **A9, A10** | Angiopoietin-1 | **D19, D20** | IL1ra | **G21, G22** | MIP3$\beta$ |
| **A11, A12** | Angiopoietin-2 | **D21, D22** | IL2 | **G23, G24** | MMP9 |

| | | | | | |
|---|---|---|---|---|---|
| **A13, A14** | BAFF | **D23, D24** | IL3 | **H1, H2** | Myeloperoxidase |
| **A15, A16** | BDNF | **E1, E2** | IL4 | **H3, H4** | Osteopontin (OPN) |
| **A17, A18** | Complement component C5/C5a | **E3, E4** | IL5 | **H5, H6** | PDGF-AA |
| **A19, A20** | CD14 | **E5, E6** | IL6 | **H7, H8** | PDGF-AB/BB |
| **A21, A22** | CD30 | **E7, E8** | IL8 | **H9, H10** | Pentraxin 3 |
| **A23, A24** | Reference Spots | **E9, E10** | IL10 | **H11, H12** | PF4 |
| **B3, B4** | CD40 ligand | **E11, E12** | IL11 | **H13, H14** | RAGE |
| **B5, B6** | Chitinase 3-like 1 | **E13, E14** | IL12 p70 | **H15, H16** | RANTES |
| **B7, B8** | Complement factor D | **E15, E16** | IL13 | **H17, H18** | RBP4 |
| **B9, B10** | C-reactive protein | **E17, E18** | IL15 | **H19, H20** | Relaxin 2 |
| **B11, B12** | Crypto-1 | **E19, E20** | IL16 | **H21, H22** | Resistin |
| **B13, B14** | Cystatin C | **E21, E22** | IL17A | **H23, H24** | SDF1α |
| **B15, B16** | DKK-1 | **E23, E24** | IL18Bpa | **I1, I2** | SERPINE1 |
| **B17, B18** | DPPIV | **F1, F2** | IL19 | **I3, I4** | SHBG |
| **B19, B20** | EGF | **F3, F4** | IL22 | **I5, I6** | ST2 |
| **B21, B22** | EMMPRIN | **F5, F6** | IL23 | **I7, I8** | TARC |
| **C3, C4** | ENA-78 | **F7, F8** | IL24 | **I9, I10** | TFF3 |
| **C5, C6** | Endolgin | **F9, F10** | IL27 | **I11, I12** | TfR |
| **C7, C8** | FAS ligand | **F11, F12** | IL31 | **I13, I14** | TGFα |

| | | | | | |
|---|---|---|---|---|---|
| **C9, C10** | FGFb | **F13, F14** | IL32 | **I15, I16** | Thrombospondin 1 |
| **C11, C12** | FGF7 | **F15, F16** | IL33 | **I17, I18** | TNFα |
| **C13, C14** | FGF19 | **F17, F18** | IL34 | **I19, I20** | uPAR |
| **C15, C16** | FLT3 ligand | **F19, F20** | IP10 | **I21, I22** | VEGF |
| **C17, C18** | G-CSF | **F21, F22** | I-TAC | **I23, I24** | Reference Spots |
| **C19, C20** | GDF15 | **F23, F24** | Kallikrein 3 | **J1, J2** | Reference Spots |
| **C21, C22** | GM-CSF | **G1, G2** | Leptin | **J3, J4** | Vitamin D BP |
| **D1, D2** | GROα | **G3, G4** | LIF | **J5, J6** | CD31 |
| **D3, D4** | Growth Hormone | **G5, G6** | Lipocalin 2 | **J7, J8** | TIM3 |
| **D5, D6** | HGF | **G7, G8** | MCP1 | **J9, J10** | VCAM1 |
| **D7, D8** | ICAM-1 | **G9, G10** | MCP3 | **J23, J24** | Negative Control |
| **D9, D10** | IFN-γ | **G11, G12** | M-CSF | | |

**Table S3.** Human cancer-related proteins tested in dot blot assay.

| **Spot Location** | **Protein** | **Spot Location** | **Protein** | **Spot Location** | **Protein** |
|---|---|---|---|---|---|
| **A1, A2** | Reference Spots | **C15, C16** | HER2 | **F5, F6** | MMP2 |
| **A3, A4** | α-Fetoprotein | **C17, C18** | HER3 | **F7, F8** | MMP3 |
| **A5, A6** | Amphiregulin | **C19, C20** | HER4 | **F9, F10** | MMP9 |

| | | | | | |
|---|---|---|---|---|---|
| **A7, A8** | Angiopoietin-1 | **C21, C22** | FGFb | **F11, F12** | MST1 |
| **A9, A10** | Angiopoietin-like 4 | **C23, C24** | - | **F13, F14** | MUC1 |
| **A11, A12** | ENPP2 | **D1, D2** | MFH1 | **F15, F16** | Nectin 4 |
| **A13, A14** | AXL | **D3, D4** | FKHR | **F17, F18** | Osteopontin |
| **A15, A16** | BCL2L1 | **D5, D6** | Galectin 3 (GAL3) | **F19, F20** | TP27/KIP1 |
| **A17, A18** | CA125 | **D7, D8** | GM-CSF | **F21, F22** | TP53 |
| **A19, A20** | E-Cadherin | **D9, D10** | HCG | **F23, F24** | PDGF-AA |
| **A21, A22** | VE-Cadherin | **D11, D12** | HGF R | **G1, G2** | CD31 |
| **A23, A24** | Reference Spots | **D13, D14** | HIF1$\alpha$ | **G3, G4** | Progesteron R |
| **B1, B2** | - | **D15, D16** | HNF3$\beta$ | **G5, G6** | Progranulin |
| **B3, B4** | CAPG | **D17, D18** | HO-1 | **G7, G8** | Prolactin |
| **B5, B6** | Carbonic Anhydrase IX | **D19, D20** | ICAM-1 | **G9, G10** | Prostasin |
| **B7, B8** | Cathepsin B | **D21, D22** | IL2ra | **G11, G12** | E-Selectin |
| **B9, B10** | Cathepsin D | **D23, D24** | IL6 | **G13, G14** | SERPINB5 (Maspin) |
| **B11, B12** | Cathepsin S | **E1, E2** | IL8 | **G15, G16** | SERPINE1 (PAI-1) |
| **B13, B14** | CEACAM-5 | **E3, E4** | IL18 BPa | **G17, G18** | SNAIL |

| | | | | | |
|---|---|---|---|---|---|
| **B15, B16** | Decorin | **E5, E6** | Kallikrein 3 | **G19, G20** | SPARC |
| **B17, B18** | DKK1 | **E7, E8** | Kallikrein 5 | **G21, G22** | Survivin |
| **B19, B20** | DLL1 | **E9, E10** | Kallikrein 6 | **G23, G24** | Tenascin C |
| **B21, B22** | HER1 | **E11, E12** | Leptin | **H1, H2** | Thrombospondin 1 |
| **B23, B24** | - | **E13, E14** | Lumican | **H3, H4** | TIE2 |
| **C1, C2** | - | **E15, E16** | CCL2/MCP1 | **H5, H6** | u-Plasminogen activator (uPA |
| **C3, C4** | Endoglin | **E17, E18** | CCL8/MCP2 | **H7, H8** | VCAM-1 |
| **C5, C6** | Endostatin | **E19, E20** | CCL7/MCP3 | **H9, H10** | VEGF |
| **C7, C8** | Enolase 2 | **E21, E22** | M-CSF | **H11, H12** | Vimentin |
| **C9, C10** | eNOS/NOS3 | **E23, E24** | Mesothelin | **I1, I2** | Reference Spots |
| **C11, C12** | EpCAM | **F1, F2** | CCL3/MIP1α | **I23, I24** | Negative Control |
| **C13, C14** | ERα | **F3, F4** | CCL20/MIP3α | | |
